# Characterization and identification of long non-coding RNAs based on feature relationship

**DOI:** 10.1101/327882

**Authors:** Guangyu Wang, Hongyan Yin, Boyang Li, Chunlei Yu, Fan Wang, Xingjian Xu, Jiabao Cao, Yiming Bao, Liguo Wang, Amir A. Abbasi, Vladimir B. Bajic, Lina Ma, Zhang Zhang

**Affiliations:** CAS Key Laboratory of Genome Sciences and Information, Beijing Institute of Genomics, Chinese Academy of Sciences, Beijing 100101, China; BIG Data Center, Beijing Institute of Genomics, Chinese Academy of Sciences, Beijing 100101, China; University of Chinese Academy of Sciences, Beijing 100049, China; Department of Biostatistics, Yale School of Public Health, New Haven, Connecticut 06520, United States; Division of Biomedical Statistics and Informatics, Mayo Clinic College of Medicine, Rochester, Minnesota 55905, United States; National Center for Bioinformatics, Programme of Comparative and Evolutionary Genomics, Faculty of Biological Sciences, Quaid-i-Azam University, Islamabad 45320, Pakistan; King Abdullah University of Science and Technology (KAUST), Computational Bioscience Research Center (CBRC), Computer, Electrical and Mathematical Sciences and Engineering Division (CEMSE), Thuwal 23955-6900, Kingdom of Saudi Arabia

**Keywords:** lncRNA, coding potential, feature relationship, ORF length, GC content

## Abstract

The significance of long non-coding RNAs (lncRNAs) in many biological processes and diseases has gained intense interests over the past several years. However, computational identification of lncRNAs in a wide range of species remains challenging; it requires prior knowledge of well-established sequences and annotations or species-specific training data, but the reality is that only a limited number of species have high-quality sequences and annotations. Here we first characterize lncRNAs by contrast to protein-coding RNAs based on feature relationship and find that the feature relationship between ORF (open reading frame) length and GC content presents universally substantial divergence in lncRNAs and protein-coding RNAs, as observed in a broad variety of species. Based on the feature relationship, accordingly, we further present LGC, a novel algorithm for identifying lncRNAs that is able to accurately distinguish lncRNAs from protein-coding RNAs in a cross-species manner without any prior knowledge. As validated on large-scale empirical datasets, comparative results show that LGC outperforms existing algorithms by achieving higher accuracy, well-balanced sensitivity and specificity, and is robustly effective (>90% accuracy) in discriminating lncRNAs from protein-coding RNAs across diverse species that range from plants to mammals. To our knowledge, this study, for the first time, differentially characterizes lncRNAs and protein-coding RNAs based on feature relationship, which is further applied in computational identification of lncRNAs. Taken together, our study represents a significant advance in characterization and identification of lncRNAs and LGC thus bears broad potential utility for computational analysis of lncRNAs in a wide range of species.

## INTRODUCTION

Long non-coding RNAs (lncRNAs) are prevalently expressed in a large number of organisms (1-5). Evidence has accumulated that lncRNAs play vital roles in biological processes including transcriptional regulation, post-transcriptional interference, translational control (6-9) and are implicated in the development of a variety of human diseases (10-14). Although the rapid advancement in DNA sequencing technologies has led to an exponential increase in the number of lncRNAs (11,15,16), lncRNAs are often tissue/cell-specific (13,17,18) and lineage/species-specific (18,19) and thus a large number of novel lncRNAs are yet to be discovered. Experimental approaches (such as ribosome profiling and mass spectrometry) for coding potential detection could provide the most direct evidence but are very time-consuming and expensive yet with limited throughput. Therefore, computational approaches are in great demand for better characterizing the landscape of lncRNAs and identifying lncRNAs in a wide variety of species.

Over the past few years, several computational algorithms have been proposed to identify lncRNAs, which fall roughly into two classes: alignment-based algorithms (20-26) and alignment-free algorithms (27-29). Representative alignment-based algorithms include CPC (Coding Potential Calculator) (21), PhyloCSF (Phylogenetic Codon Substitution Frequencies) (22), and COME (coding potential calculation tool based on multiple features) (26). To distinguish lncRNAs from protein-coding transcripts, specifically, CPC uses sequence alignments against known proteins (21), PhyloCSF relies on multiple alignments of sequences from closely related species (22), and COME integrates multiple sequence-derived and experiment-based features (including DNA conservation, protein conservation, RNA structure conservation, GC content, expression, histone methylation) (26). Clearly, alignment-based algorithms are limited by the completeness of known proteins and the accuracy of DNA alignments and some of them are highly dependent on experiment-based features. Most importantly, they are incapable of identifying lncRNAs that are lineage/species-specific (18,19) and become unreliable when no high-quality genome annotation is available. Additionally, alignment-based algorithms require prior sequence alignments and thus are exceedingly time-consuming, especially when more and more known sequences become available.

In contrast, alignment-free algorithms do not need any alignment but require high-quality protein-coding RNAs and lncRNAs as training data (27-30). Representative algorithms include CPAT (Coding-Potential Assessment Tool) (28), CNCI (Coding-Non-Coding Index) (27) and PLEK (predictor of long non-coding RNAs and messenger RNAs based on an improved *k*-mer scheme) (29). However, most of these algorithms are species-specific. These algorithms become unreliable when they are trained on data from one species and applied to data from another species (24,27,29). Moreover, alignment-free algorithms are heavily dependent on high-quality training data, but in reality, many species have low-quality or even no annotations, especially for newly sequenced species. It is reported that there are ∼8.7 million eukaryotic species on Earth and ∼90% species’ genomes are still waiting to be deciphered (31). Therefore, it is desirable to develop a more robust and effective algorithm that is able to accurately distinguish lncRNAs from protein-coding RNAs without the need of any prior information on alignment or training.

It would be straightforward to classify lncRNAs and protein-coding RNAs by taking account of sequence features. Although sequence features have already been factored in existing algorithms, for instance, ORF (open reading frame) length and coverage (20,21,24,25,28), sequence similarity and conservation (20-25), nucleotide composition and codon usage (20,24,26-29), existing algorithms regard sequence features as independent variables and do not consider their potential biological relationship. Here we characterize lncRNAs by contrast to protein-coding RNAs based on a feature relationship between ORF length and GC content. As this feature relationship presents universally substantial divergence between lncRNAs and protein-coding RNAs as observed in a wide variety of species, we further propose LGC (ORF Length and GC content), a novel algorithm for robust and effective discrimination of lncRNAs from protein-coding RNAs. As testified on large-scale empirical datasets, LGC represents a significant advance over existing algorithms by identifying lncRNAs in a wide range of species not only effectively but also robustly. To our knowledge, this is the first to differentially characterize lncRNAs and protein-coding RNAs based on feature relationship, which is further applicable and effective in accurate identification of lncRNAs.

## METHODS AND MATERIALS

### Modeling the Relationship between ORF Length and GC Content

It has been reported that in an ORF with random distribution of nucleotide, the expected ORF length increases with its GC content (32). More specifically, in an unbiased sequence, where the frequency of adenine is equal to that of thymine, and the frequency of guanine is equal to that of cytosine, i.e., *P*_*A*_=*P*_*T*_ and *P*_*G*_=*P*_*C*_, the probability of observing a stop-codon (*f*) as reported in (32), can be expressed as

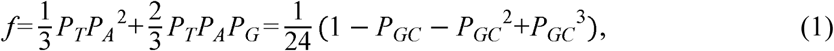

where *P*_*GC*_ is the GC content and equals to *P*_*G*_ + *P*_*C*_. However, the presence of intron (33), mutation pressure on different exons (34), and selection against cytosine (C) usage (35) will modulate the relationship between ORF length and GC content, so that the stop-codon probability can hardly be inferred from Eq. (1). Therefore, we compose a more flexible equation (Eq. (2)), which utilizes four parameters (*a*_0_, *a*_1_, *a*_2_, *a*_3_) to reflect the different relationship between GC content and stop-codon probability (*f*) in different genomic background

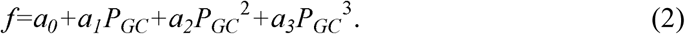

Accordingly, the expected length of ORF is 3/*f*. Because of a potential bias from short sequences, we consider only ORFs longer than 100nt and thus, the expected ORF length (E(*l*)) can be expressed as

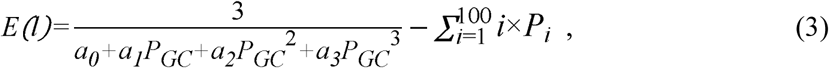

where *P*_*i*_ is the frequency of ORFs with the length of *i* nt (ranging from 1 to 100). Then we use polynomial function of GC content to approximate 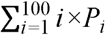.

To investigate the relationship between ORF length and GC content, we choose the top three longest ORFs (longer than 100nt) for each sequence and divide ORFs into 100 groups based on their GC contents. Mean estimates of ORF length and GC content are used to estimate the parameters of Eq. (3) by the least square method. Root mean square error (RMSE) is used as the criterion function for fitting the model of the expected ORF length from Eq. (3) for both protein-coding RNAs and lncRNAs (Table 1).

**Table 1.** Parameters for species-specific model

| Species | Protein-coding RNA |  |  |  |  | LncRNA |  |  |  |  |
| --- | --- | --- | --- | --- | --- | --- | --- | --- | --- | --- |
| | $a_0$ | $a_1$ | $a_2$ | $a_3$ | RMSE* | $a_0$ | $a_1$ | $a_2$ | $a_3$ | RMSE* |
| *H. sapiens* | 0.0247 | -0.1109 | 0.1781 | -0.0933 | 106.14 | 0.0018 | 0.0256 | -0.0577 | 0.0356 | 24.94 |
| *M. musculus* | 0.0295 | -0.1241 | 0.1773 | -0.0776 | 109.30 | 0.0064 | -0.0035 | 0.0024 | -0.0043 | 21.36 |
| *D. rerio* | 0.0499 | -0.2116 | 0.2772 | -0.0941 | 188.41 | 0.0200 | -0.0705 | 0.0919 | -0.0315 | 53.81 |
| *C. elegans* | 0.0707 | -0.4205 | 0.8338 | -0.5340 | 136.26 | 0.0067 | -0.0069 | 0.0084 | -0.0001 | 16.97 |
| *O. sativa* | 0.0246 | -0.1130 | 0.1803 | -0.0933 | 114.11 | 0.0177 | -0.0773 | 0.1568 | -0.1033 | 27.48 |
| *S. lycopersicum* | 0.0649 | -0.3437 | 0.5536 | -0.2249 | 177.46 | -0.0478 | 0.3571 | -0.7745 | 0.5419 | 66.07 |

\* RMSE: root-mean-square error; see Eq. (2) for more information on parameters ( $a_0$ , $a_1$ , $a_2$ , $a_3$ ).

### Maximum Likelihood Estimation of Coding Potential

Protein-coding RNAs and lncRNAs are used to fit Eq. (3) to estimate parameters *a*_0_, *a*_1_, *a*_2_ and *a*_3_, and these estimates are then applied to Eq. (2), from which the probability of stop codon can be derived. For any given transcript that has *n* sense codons, its coding potential score (*L*) can be estimated by the maximum likelihood method through calculating the log likelihood ratio based on Eq. (4)

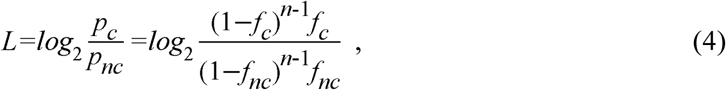

where *p*_*c*_ is the probability of ORF in coding sequence, *p*_*nc*_ is the probability of ORF in non-coding sequence, *f*_*c*_ is the probability of finding a stop codon in coding sequence, and *f*_*nc*_ is the probability of finding a stop codon in non-coding sequence. *L*>0 indicates it is a protein-coding RNA and *L*<0 indicates that it is a non-coding RNA. Symbols used in calculating coding potential score are listed in Table 2.

**Table 2.**
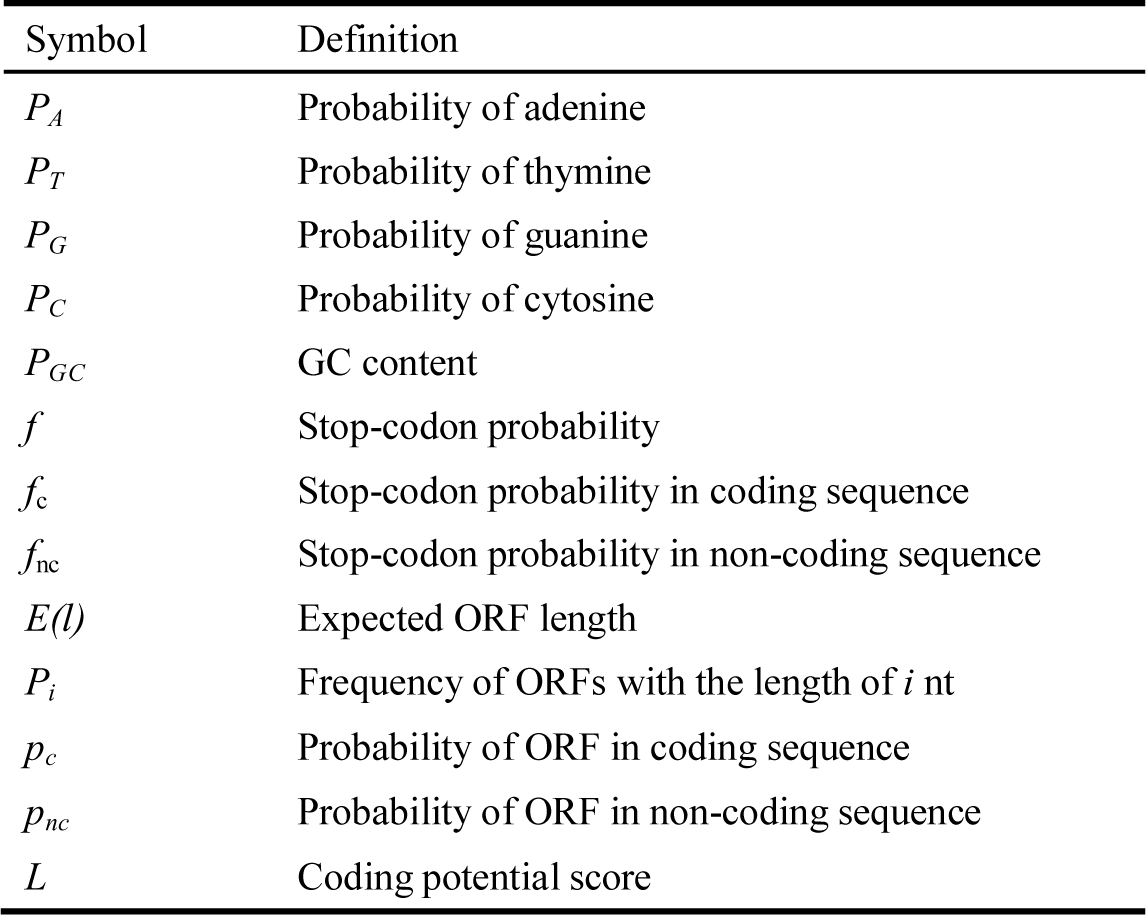
Symbols used in calculating coding potential score

| Symbol | Definition |
| --- | --- |
| $P_A$ | Probability of adenine |
| $P_T$ | Probability of thymine |
| $P_G$ | Probability of guanine |
| $P_C$ | Probability of cytosine |
| $P_{GC}$ | GC content |
| $f$ | Stop-codon probability |
| $f_c$ | Stop-codon probability in coding sequence |
| $f_{nc}$ | Stop-codon probability in non-coding sequence |
| $E(l)$ | Expected ORF length |
| $P_i$ | Frequency of ORFs with the length of $i$ nt |
| $p_c$ | Probability of ORF in coding sequence |
| $p_{nc}$ | Probability of ORF in non-coding sequence |
| $L$ | Coding potential score |

### Performance Evaluation of LGC

Protein-coding RNAs (38,811 transcripts) and lncRNAs (27,669 transcripts) of human (Table S1) are used to build LGC. Ten-fold cross-validation shows that LGC achieves very high accuracy on human data, with an AUC of 0.981 (Figure S1). LGC is evaluated by comparison with several existing popular algorithms, including CPC (21), CPAT (28), CNCI (27), and PLEK (29). LGC, CPC, and PLEK can be used in a cross-species manner that do not require any training or specific model. CNCI is also used in a cross-species manner, but uses two specific models, namely, “*ve*” and “*pl*”, to identify lncRNAs in animals (human, mouse, zebrafish, worm) and plants (rice and tomato), respectively. Contrastingly, CPAT uses species-specific training data to build specific models. Specifically, it adopts species-specific logistic regression models to calculate coding probability and sets different cutoffs, viz., 0.36 for human, 0.44 for mouse, 0.38 for zebrafish, and 0.39 for worm. Due to the lack of a prebuilt model for plant, the logistic regression model of human is additionally applied to rice and tomato during performance comparison. We compare LGC with algorithms that can be used in a cross-species manner or adopt specific models. All datasets used for comparisons are summarized into Table S1. To reduce any bias from unequal sampling size of lncRNAs and protein-coding RNAs, we randomly select protein-coding RNAs with the equal number of lncRNAs.

To compare the performance of different algorithms in distinguishing lncRNAs from protein-coding RNAs, protein-coding RNAs and lncRNAs are denoted as positive and negative samples, respectively. As a result, *accuracy, sensitivity*, and *specificity* can be estimated according to Eqs. (5-7), which take account of true positive (*TP*), true negative (*TN*), false positive (*FP*) and false negative (*FN*) predictions.

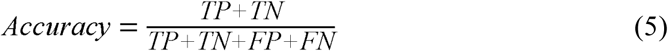

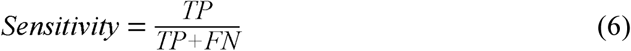

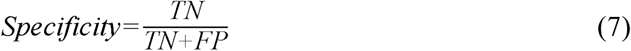

### Data Collection

A total of six representative organisms are used in this study, including two mammals (human and mouse), one vertebrate (zebrafish), one invertebrate (worm) and two plants (rice and tomato). Protein-coding RNAs for human and mouse are both collected from NCBI RefSeq (36) and their corresponding lncRNAs are obtained from Gencode version 22 (37) and Gencode version M7 (38), respectively. For the remaining organisms, both protein-coding RNAs and ncRNAs are downloaded from Ensembl (39). To obtain lncRNAs, ncRNAs shorter than 200nt are excluded. All detailed information of these datasets is summarized in Table S1.

### Availability

The package of LGC can be downloaded for academic use only at BioCode (a source code archive for bioinformatics software tools; http://bigd.big.ac.cn/biocode) in the BIG Data Center (40), with accession number BT000004. In addition, a web server is publicly available at http://bigd.big.ac.cn/lgc/calculator.

## RESULTS AND DISCUSSION

### Characterization of Protein-coding RNAs and LncRNAs Based on Feature Relationship

It is extensively documented that in protein-coding sequences ORF length is dominantly determined by GC content, since base composition of translational stop codons (TAG, TAA, and TGA) is biased toward low GC content (32,34,35). If a sequence is AT-rich, it is most likely that stop codons would appear earlier, resulting in a shorter ORF; conversely, a GC-rich sequence tends to have longer ORF because it is less likely to have stop codons earlier (32). Considering that protein-coding RNAs differ from lncRNAs in possessing significantly longer ORFs (21,24,25,28), it is possible that protein-coding RNAs and lncRNAs may present different relationships between GC content and ORF length. Of course, ORF length, as one of the important features, has been widely used by the existing algorithms in coding potential prediction (20,21,24,25,28). However, the two features—ORF length and GC content—are often regarded as independent, and their relationship has not been well characterized in protein-coding RNAs and lncRNAs. Therefore, we model the relationship between ORF length and GC content and hypothesize that this relationship can be used to differentially characterize protein-coding RNAs and lncRNAs.

To test the hypothesis, we collect protein-coding RNAs and lncRNAs from six representative organisms (Table S1) and examine their corresponding relationships between ORF length and GC content (based on Eq. (3); see Materials and Methods). Consistent with our expectations, protein-coding RNAs and lncRNAs present strikingly different relationships in all investigated organisms (Figure 1). An obvious inverted V-shape curve is observed in protein-coding RNAs, that is, ORF length increases with GC content for low-GC genes, while decreases for high-GC genes. This is well consistent with pervious findings that selection against cytosine usage (prone to mutation to T/U; e.g. CAR to TAR and CGA to TGA) (35) in GC-rich genes may contribute to negative correlation between GC content and ORF length. Compared to protein-coding RNAs, contrastingly, curves are extremely flat in lncRNAs. Overall, these results show that protein-coding RNAs and lncRNAs exhibit significant and universal heterogeneity in the relationship between ORF **L**ength and **GC** content (**LGC** model). Thus, based on the LGC model, we further explore whether such heterogeneity can be used to effectively distinguish lncRNAs from protein-coding RNAs for a wide variety of species.

**Figure 1.**
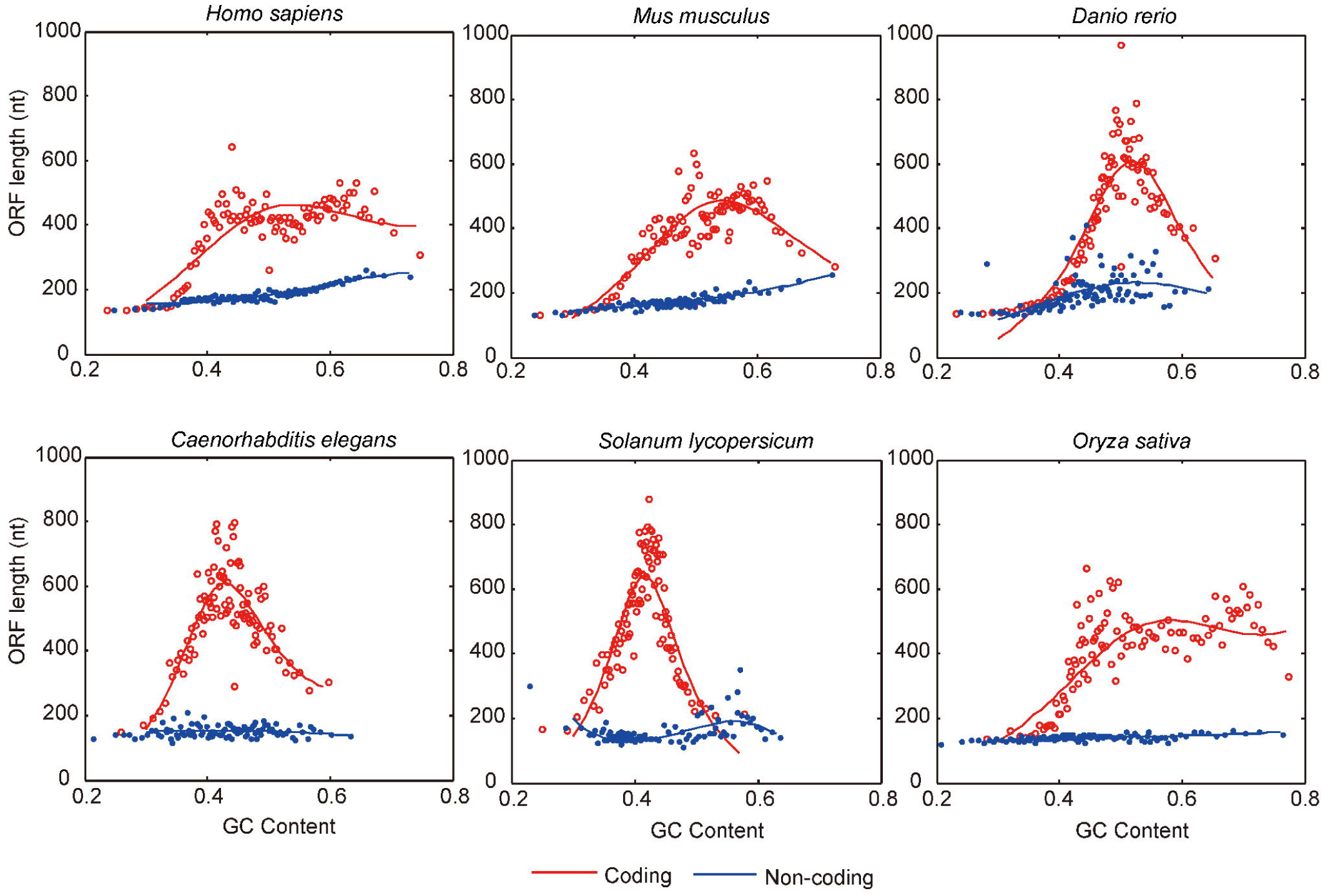
Relationship between ORF length and GC content for protein-coding RNAs (red circles) and lncRNAs (blue dots), respectively. For each transcript, the top three longest ORFs (longer than 100nt) are used. ORFs are grouped into 100 bins based on their GC contents and each dot represents the averaged estimate for each bin.

### Application of the LGC Model in LncRNA Identification

To apply the LGC model in the identification of lncRNAs, we first estimate parameters in Eqs. (2-3) (see Methods and Materials) using all lncRNAs and protein-coding RNAs for each species and build species-specific LGC (Table 1). We then employ these parameters’ estimates to calculate the coding potential score (Eq. (4)), which is an indicator to distinguish lncRNAs from protein-coding RNAs. As validated on empirical datasets from six representative species (Table 3), we find that species-specific LGC model achieves high accuracy (>0.88) in each species and performs well in the identification of both protein-coding RNAs and lncRNAs as indicated by well-balanced sensitivities and specificities. These results suggest that the LGC model is indeed applicable for identifying lncRNAs in a wide range of species.

**Table 3.** Performance of LGC based on species-specific model and human model

| Species | Species-specific model |  |  | Human model |  |  |
| --- | --- | --- | --- | --- | --- | --- |
|  | Accuracy | Sensitivity | Specificity | Accuracy | Sensitivity | Specificity |
| *H. sapiens* | 0.942 | 0.964 | 0.921 | 0.942 | 0.964 | 0.921 |
| *M. musculus* | 0.956 | 0.948 | 0.965 | **0.965** | **0.960** | **0.969** |
| *D. rerio* | 0.884 | 0.875 | **0.893** | **0.911** | **0.939** | 0.882 |
| *C. elegans* | 0.933 | 0.868 | 0.996 | **0.952** | **0.908** | **0.996** |
| *O. sativa* | 0.929 | **0.915** | 0.944 | **0.931** | 0.909 | **0.953** |
| *S. lycopersicum* | 0.899 | 0.805 | 0.993 | **0.920** | **0.844** | **0.995** |

\* The species-specific model indicates that LGC is built based on species-specific protein-coding RNAs and lncRNAs. The human model indicates that LGC is built based only on human protein-coding RNAs and lncRNAs. These models are compared in terms of accuracy, sensitivity, and specificity, where numbers in bold represent the better performance.

To further test the universality of the LGC model across different species, we compare the performances of species-specific LGC model against the LGC model built based merely on human data (whose quality is believed to be relatively higher). Strikingly, the LGC model based on human data overall shows better performances than the species-specific LGC models in terms of accuracy, specificity, and sensitivity (Table 3); it achieves higher accuracies (>0.91) in all six organisms, with both sensitivities and specificities greater than 0.84. Although the LGC model is built bases on human data, high accuracy is achieved not only for mammals and vertebrates but also for invertebrates and plants. This result, which is most likely caused by both larger-quantity and higher-quality of human data, suggests that the LGC model is universally applicable, guaranteeing that the LGC model can be used in a cross-species manner without requiring species-specific data.

### Effective Discrimination of LncRNAs from Protein-coding RNAs

To test the effectiveness of the LGC model and to examine its performance in discriminating lncRNAs from protein-coding RNAs, we further evaluate it on data from six representative organisms by comparison with several popular algorithms including CPC (21), CNCI (27), and PLEK (29) that can be used in a cross-species manner (see Methods and Materials). In what follows, the LGC model built on human data (as detailed above), is used in all comparisons.

Comparative evaluations regarding accuracy, specificity, and sensitivity show that LGC outperforms existing algorithms, significantly and robustly across different species (Figure 2). Specifically, LGC overall achieves higher accuracies for all six organisms (higher than 0.91); it outperforms CNCI and PLEK in non-human species, and CPC in human, mouse, zebrafish and rice. Considering the average accuracy over all six species (Table 4), LGC obtains the highest average accuracy (0.937) compared with CPC (0.903), CNCI (0.895) and PLEK (0.822). Moreover, LGC yields better average specificity of 0.953 across all six species than the other algorithms (Table 4); it outperforms CNCI in zebrafish and rice, PLEK in mouse, zebrafish and rice, and CPC in human, mouse, zebrafish and rice (and performs comparably in the remaining cases) (Figure 2). Regarding sensitivity, LGC achieves the average sensitivity at 0.921 (just follows CPC at 0.981), better than CNCI at 0.848, and PLEK at 0.756 (Table 4); it outperforms CPC in mouse, CNCI in human, worm and tomato, and PLEK in mouse, worm, tomato and rice (Figure 2).

**Table 4.** Estimates of accuracy, sensitivity, and specificity averaged over six representative organisms

| Algorithm | LGC | CNCI | CPAT | CPC | PLEK |
| --- | --- | --- | --- | --- | --- |
| Accuracy | **0.937** | 0.895 | 0.925 | 0.903 | 0.822 |
| Sensitivity | 0.921 | 0.848 | 0.910 | **0.981** | 0.756 |
| Specificity | **0.953** | 0.941 | 0.940 | 0.823 | 0.936 |

\* Numbers in bold represent the better performance.

**Figure 2.**
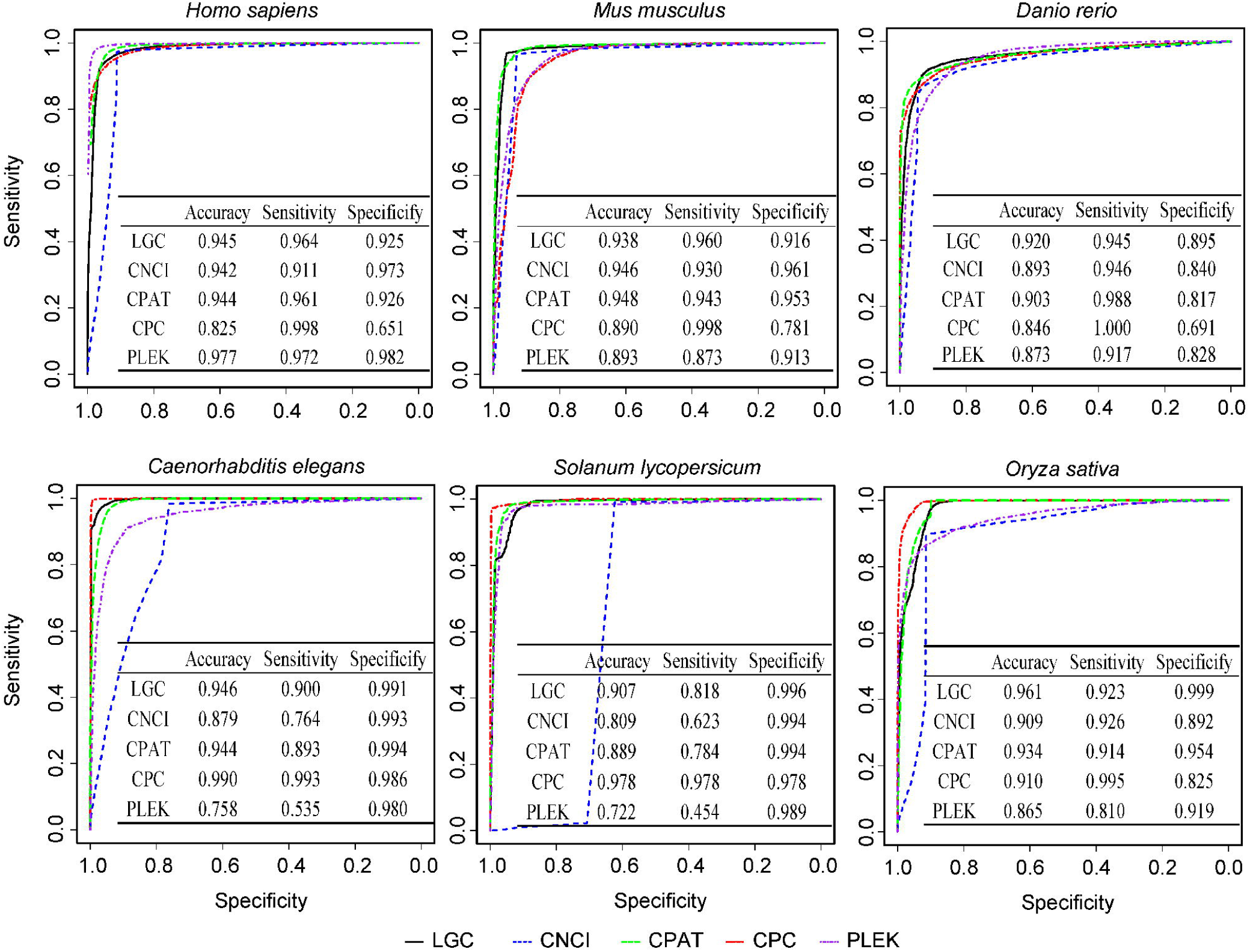
Performances of LGC, CNCI, CPAT, CPC, and PLEK. LGC, CPC, CNCI, and PLEK can be used in a cross-species manner, while CPAT uses specific models and cutoffs for different species (see Methods and Materials).

Strikingly, LGC provides well-balanced sensitivity and specificity (both higher than 84%), which is consistently observed for all examined species (Figure 2). Contrary to this, existing algorithms show poor balance between sensitivity and specificity; CPC yields extremely unbalanced sensitivity and specificity (for instance, 0.998 and 0.651 in human, respectively), CNCI presents sensitivity at 0.761 and specificity at 0.984 in worm (consistent with the previous study in (27)), and PLEK exhibits sensitivity at 0.527 and specificity at 0.980 in worm. Taken together, these results clearly show that LGC achieves a good balance between sensitivity and specificity and is capable of discriminating lncRNAs from protein-coding RNAs more accurately than the existing algorithms.

To further evaluate the performance of LGC, we also compare it with CPAT (28), which requires appropriate training to build specific models with different cutoff values (see details in Methods and Materials). Albeit CPAT uses species-specific models, we find that LGC overall performs better than CPAT (Figure 2 and Table 4). Specifically, it performs comparably with CPAT in human and outperforms CPAT in non-human species (Figure 2). Although CPAT builds species-specific models for human, mouse, zebrafish, and fly, it does not perform well as expected. In zebrafish, CPAT shows poor balance between sensitivity (0.988) and specificity (0.817), whereas LGC yields sensitivity at 0.939 and specificity at 0.882. This may because that training data of different models show unequal qualities, and the robust performance of human, mouse, and fly models of CPAT are attributable to the high quality of training data sets. Also, it is noted that species-specific algorithms have significant limitation in application. As no prebuilt models are available for plants, we apply human model of CPAT to tomato and rice. However, the human model of CPAT presents unbalanced sensitivity (at 0.783) and specificity (at 0.994) in tomato, whereas LGC yields sensitivity at 0.844 and specificity at 0.995. Given that CPAT is heavily dependent on high-quality training data and many species presently may still have low-quality or even no training data, LGC bears broad utility for computational analysis of lncRNAs in a wide range of species.

### Robustness in a Wide Diversity of Species

To further examine the robustness of LGC for a wider diversity of species, we set up a more comprehensive dataset by collecting all curated protein-coding RNAs (accession prefixed with NM) and ncRNAs (accession prefixed with NR) from NCBI RefSeq (36). All protein-coding RNAs and ncRNAs are classified into: mammals (127,903 protein-coding RNAs from 81 species and 23,644 ncRNAs from 26 species), vertebrates (53,239 protein-coding RNAs from 59 species and 2,582 ncRNAs from 9 species), invertebrates (68,229 protein-coding RNAs from 42 species and 29,527 ncRNAs from 11 species) and plants (97,119 protein-coding RNAs from 34 species and 1,795 ncRNAs from 10 species).

We test the performance of LGC on this more comprehensive dataset derived from a larger number of species and compare it against existing algorithms that can be used in a cross-species manner without requiring any species-specific training or model. Accordingly, only PLEK, albeit built on human data, can be used for a wide range of species (29), whereas other algorithms are unsuitable for this comparison (as CNCI is limited to two specific models, namely, ‘ve’ for vertebrates, and ‘pl’ for plants (27), CPC depends on sequence alignments against known proteins (21), which are completely identical to the dataset obtained from NCBI RefSeq (36)). Comparative results show that in general LGC performs more stable and achieves higher accuracy (>0.9 for most datasets) in the identification of both protein-coding and ncRNAs (Figure 3). By contrast, PLEK, based on a *k*-mer scheme and a support vector machine (SVM) algorithm (29), performs poorly and shows an obvious imbalance in its ability to identify both protein-coding and non-coding RNAs for all investigated cases (Figure 3). In addition, PLEK presents unstable varied performances in plants, vertebrates, invertebrates and mammals, whereas LGC achieves robust higher accuracies in almost all datasets (Figure 3, Tables S2 and S3). Collectively, these results indicate that LGC is robust in accurately discriminating lncRNAs from protein-coding RNAs in a wide variety of species.

**Figure 3.**
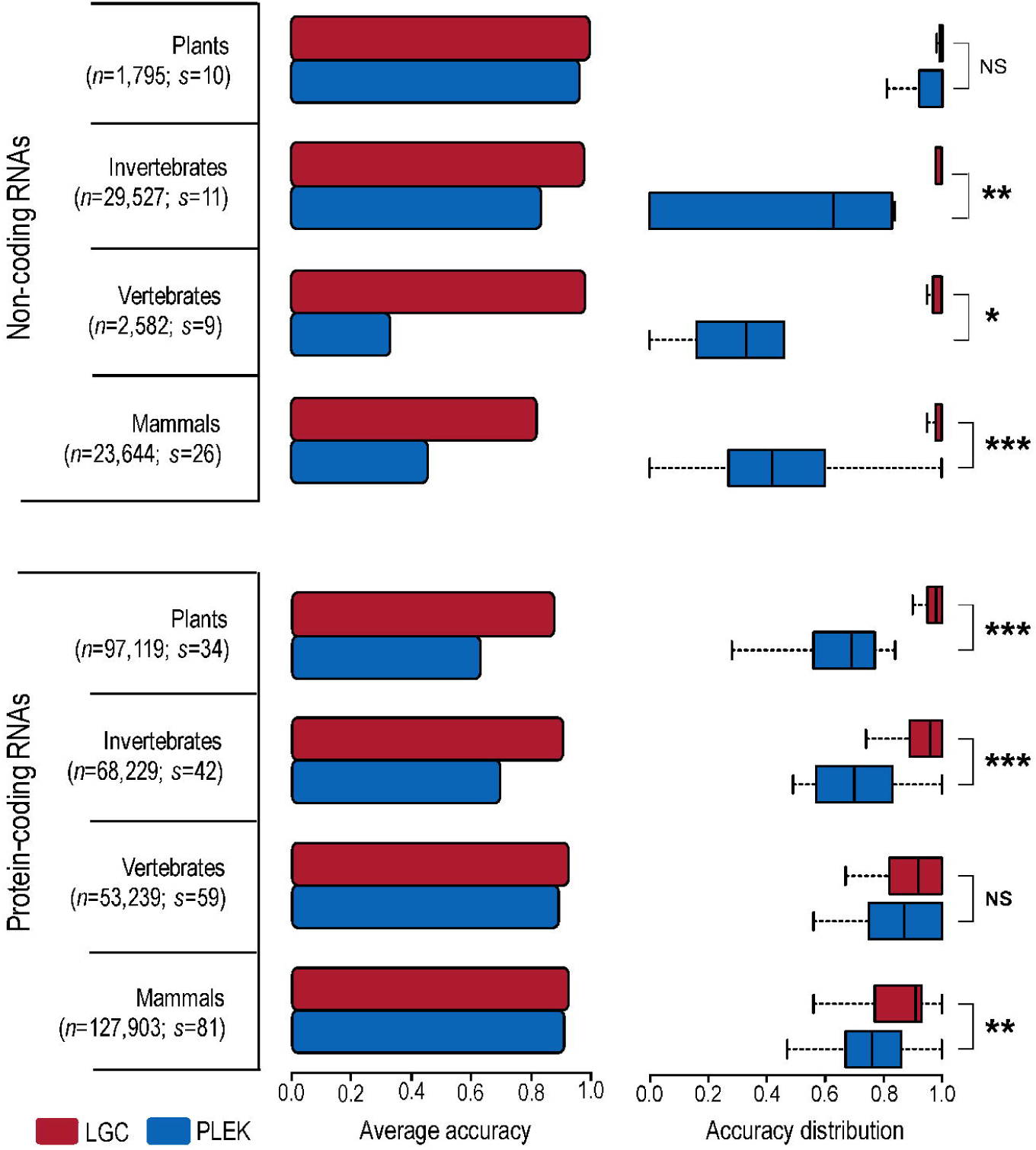
Accuracy of LGC and PLEK on protein-coding RNAs and non-coding RNAs from NCBI RefSeq. The number of sequences (*n*) as well as the number of species (*s*) is labeled. The boxes depict data between the 25th and 75th percentiles with central horizontal lines representing the median values. The Wilcoxon test is used to evaluate the significance level when comparing the accuracy between LGC and PLEK, and *P*-value is indicated by NS (Not Significant) > 0.05, ‘*’ ≤ 0.05, ‘**’ ≤ 10^−3^, and ‘***’ ≤ 10^−5^. Comparison results for each species are listed in Table S2 (non-coding RNA) and Table S3 (protein-coding RNA).

## CONCLUSION

To our knowledge, our study is the first to differentially characterize lncRNAs and protein-coding RNAs based on a feature relationship between ORF length and GC content, on the grounds that lncRNAs and protein-coding RNAs present considerable divergence in terms of this relationship, which is consistently and universally detected in a wide range of species. Hence, we further present LGC, a novel algorithm to discriminate lncRNAs from protein-coding RNAs based on this feature relationship. As demonstrated in multiple empirical datasets across a wide diversity of species, LGC is superior to existing algorithms by achieving higher accuracy and well-balanced sensitivity and specificity. In addition, LGC is able to accurately and robustly distinguish lncRNAs from protein-coding RNAs in a cross-species manner without the need for species-specific adjustments. Overall, LGC represents a simple, robust and powerful algorithm for characterization and identification of lncRNAs in a wide range of species, providing a significant advance for computational analysis of lncRNAs.

## ACKNOWLEDGEMENT

We thank Shuhui Song, Lili Hao, and Shixiang Sun for valuable comments on this work.

## AUTHOR CONTRIBUTIONS

LM and ZZ conceived and designed the project. Model development: GW and LM; Program and web server coding: GW, LM, BL, FW, XX, JC, LW; Data analysis: LM, GW, HY, CY; Manuscript draft: LM and GW. AA, LW, VBB and YB revised the manuscript. ZZ supervised this project and revised the manuscript.

## REFERENCES

1. Carninci, P., Kasukawa, T., Katayama, S., Gough, J., Frith, M.C., Maeda, N., Oyama, R., Ravasi, T., Lenhard, B., Wells, C. et al. (2005) The transcriptional landscape of the mammalian genome. Science, 309, 1559–1563.

2. Kapranov, P., Cheng, J., Dike, S., Nix, D.A., Duttagupta, R., Willingham, A.T., Stadler, P.F., Hertel, J., Hackermuller, J., Hofacker, I.L. et al. (2007) RNA maps reveal new RNA classes and a possible function for pervasive transcription. Science, 316, 1484–1488.

3. Pennisi, E. (2010) Shining a light on the genome’s ‘dark matter’. Science, 330, 1614.

4. Djebali, S., Davis, C.A., Merkel, A., Dobin, A., Lassmann, T., Mortazavi, A., Tanzer, A., Lagarde, J., Lin, W., Schlesinger, F. et al. (2012) Landscape of transcription in human cells. Nature, 489, 101–108.

5. Liu, X., Hao, L., Li, D., Zhu, L. and Hu, S. (2015) Long non-coding RNAs and their biological roles in plants. Genomics Proteomics Bioinformatics, 13, 137–147.

6. Wilusz, J.E., Sunwoo, H. and Spector, D.L. (2009) Long noncoding RNAs: functional surprises from the RNA world. Genes Dev, 23, 1494–1504.

7. Mercer, T.R., Dinger, M.E. and Mattick, J.S. (2009) Long non-coding RNAs: insights into functions. Nat Rev Genet, 10, 155–159.

8. Rinn, J.L. and Chang, H.Y. (2012) Genome regulation by long noncoding RNAs. Annu Rev Biochem, 81, 145–166.

9. Chen, H., Du, G., Song, X. and Li, L. (2017) Non-coding Transcripts from Enhancers: New Insights into Enhancer Activity and Gene Expression Regulation. Genomics Proteomics Bioinformatics, 15, 201–207.

10. Chen, G., Wang, Z., Wang, D., Qiu, C., Liu, M., Chen, X., Zhang, Q., Yan, G. and Cui, Q. (2013) LncRNADisease: a database for long-non-coding RNA-associated diseases. Nucleic Acids Res, 41, D983–986.

11. Ma, L.N., Li, A., Zou, D., Xu, X.J., Xia, L., Yu, J., Bajic, V.B. and Zhang, Z. (2015) LncRNAWiki: harnessing community knowledge in collaborative curation of human long non-coding RNAs. Nucleic Acids Res, 43, D187–D192.

12. Salhi, A., Essack, M., Alam, T., Bajic, V.P., Ma, L., Radovanovic, A., Marchand, B., Schmeier, S., Zhang, Z. and Bajic, V.B. (2017) DES-ncRNA: A knowledgebase for exploring information about human micro and long noncoding RNAs based on literature-mining. RNA Biol, 14, 963–971.

13. Alam, T., Uludag, M., Essack, M., Salhi, A., Ashoor, H., Hanks, J.B., Kapfer, C., Mineta, K., Gojobori, T. and Bajic, V.B. (2017) FARNA: knowledgebase of inferred functions of non-coding RNA transcripts. Nucleic Acids Res, 45, 2838–2848.

14. Fang, Y. and Fullwood, M.J. (2016) Roles, Functions, and Mechanisms of Long Non-coding RNAs in Cancer. Genomics Proteomics Bioinformatics, 14, 42–54.

15. Iyer, M.K., Niknafs, Y.S., Malik, R., Singhal, U., Sahu, A., Hosono, Y., Barrette, T.R., Prensner, J.R., Evans, J.R., Zhao, S. et al. (2015) The landscape of long noncoding RNAs in the human transcriptome. Nat Genet, 47, 199–208.

16. Zhao, Y., Li, H., Fang, S., Kang, Y., Wu, W., Hao, Y., Li, Z., Bu, D., Sun, N., Zhang, M.Q. et al. (2016) NONCODE 2016: an informative and valuable data source of long non-coding RNAs. Nucleic Acids Res, 44, D203–208.

17. Cabili, M.N., Trapnell, C., Goff, L., Koziol, M., Tazon-Vega, B., Regev, A. and Rinn, J.L. (2011) Integrative annotation of human large intergenic noncoding RNAs reveals global properties and specific subclasses. Genes Dev, 25, 1915–1927.

18. Derrien, T., Johnson, R., Bussotti, G., Tanzer, A., Djebali, S., Tilgner, H., Guernec, G., Martin, D., Merkel, A., Knowles, D.G. et al. (2012) The GENCODE v7 catalog of human long noncoding RNAs: analysis of their gene structure, evolution, and expression. Genome Res, 22, 1775–1789.

19. Paralkar, V.R., Mishra, T., Luan, J., Yao, Y., Kossenkov, A.V., Anderson, S.M., Dunagin, M., Pimkin, M., Gore, M., Sun, D. et al. (2014) Lineage and species-specific long noncoding RNAs during erythro-megakaryocytic development. Blood, 123, 1927–1937.

20. Liu, J.F., Gough, J. and Rost, B. (2006) Distinguishing protein-coding from non-coding RNAs through support vector machines. Plos Genetics, 2, 529–536.

21. Kong, L., Zhang, Y., Ye, Z.Q., Liu, X.Q., Zhao, S.Q., Wei, L. and Gao, G. (2007) CPC: assess the protein-coding potential of transcripts using sequence features and support vector machine. Nucleic Acids Res, 35, W345–349.

22. Lin, M.F., Jungreis, I. and Kellis, M. (2011) PhyloCSF: a comparative genomics method to distinguish protein coding and non-coding regions. Bioinformatics, 27, i275–282.

23. Washietl, S., Findeiss, S., Muller, S.A., Kalkhof, S., von Bergen, M., Hofacker, I.L., Stadler, P.F. and Goldman, N. (2011) RNAcode: robust discrimination of coding and noncoding regions in comparative sequence data. RNA, 17, 578–594.

24. Achawanantakun, R., Chen, J., Sun, Y.N. and Zhang, Y. (2015) LncRNA-ID: Long non-coding RNA IDentification using balanced random forests. Bioinformatics, 31, 3897–3905.

25. Sun, K., Chen, X.N., Jiang, P.Y., Song, X.F., Wang, H.T. and Sun, H. (2013) iSeeRNA: identification of long intergenic non-coding RNA transcripts from transcriptome sequencing data. Bmc Genomics, 14, S7.

26. Hu, L., Xu, Z., Hu, B. and Lu, Z.J. (2016) COME: a robust coding potential calculation tool for lncRNA identification and characterization based on multiple features. Nucleic Acids Res, 45, e2.

27. Sun, L., Luo, H., Bu, D., Zhao, G., Yu, K., Zhang, C., Liu, Y., Chen, R. and Zhao, Y. (2013) Utilizing sequence intrinsic composition to classify protein-coding and long non-coding transcripts. Nucleic Acids Res, 41, e166.

28. Wang, L., Park, H.J., Dasari, S., Wang, S., Kocher, J.P. and Li, W. (2013) CPAT: Coding-Potential Assessment Tool using an alignment-free logistic regression model. Nucleic Acids Res, 41, e74.

29. Li, A., Zhang, J. and Zhou, Z. (2014) PLEK: a tool for predicting long non-coding RNAs and messenger RNAs based on an improved k-mer scheme. BMC Bioinformatics, 15, 311.

30. Alam, T., Medvedeva, Y.A., Jia, H., Brown, J.B., Lipovich, L. and Bajic, V.B. (2014) Promoter analysis reveals globally differential regulation of human long non-coding RNA and protein-coding genes. PLoS One, 9, e109443.

31. Mora, C., Tittensor, D.P., Adl, S., Simpson, A.G. and Worm, B. (2011) How many species are there on Earth and in the ocean? PLoS Biol, 9, e1001127.

32. Oliver, J.L. and Marin, A. (1996) A relationship between GC content and coding-sequence length. J Mol Evol, 43, 216–223.

33. Senapathy, P. (1986) Origin of Eukaryotic Introns - a Hypothesis, Based on Codon Distribution Statistics in Genes, and Its Implications. P Natl Acad Sci USA, 83, 2133–2137.

34. Xia, X.H., Xie, Z. and Li, W.H. (2003) Effects of GC content and mutational pressure on the lengths of exons and coding sequences. J Mol Evol, 56, 362–370.

35. Xia, X.H., Wang, H.C., Xie, Z., Carullo, M., Huang, H. and Hickey, D. (2006) Cytosine usage modulates the correlation between CDS length and CG content in prokaryotic genomes. Mol Biol Evol, 23, 1450–1454.

36. Pruitt, K.D., Tatusova, T. and Maglott, D.R. (2007) NCBI reference sequences (RefSeq): a curated non-redundant sequence database of genomes, transcripts and proteins. Nucleic Acids Res, 35, D61–65.

37. Harrow, J., Frankish, A., Gonzalez, J.M., Tapanari, E., Diekhans, M., Kokocinski, F., Aken, B.L., Barrell, D., Zadissa, A., Searle, S. et al. (2012) GENCODE: the reference human genome annotation for The ENCODE Project. Genome Res, 22, 1760–1774.

38. Mudge, J.M. and Harrow, J. (2015) Creating reference gene annotation for the mouse C57BL6/J genome assembly. Mamm Genome, 26, 366–378.

39. Cunningham, F., Amode, M.R., Barrell, D., Beal, K., Billis, K., Brent, S., Carvalho-Silva, D., Clapham, P., Coates, G., Fitzgerald, S. et al. (2015) Ensembl 2015. Nucleic Acids Res, 43, D662–669.

40. BIG Data Center Members. (2018) Database Resources of the BIG Data Center in 2018. Nucleic Acids Research, 45, D18–D24.

